# Multi-trait regressor stacking increased genomic prediction accuracy of sorghum grain composition

**DOI:** 10.1101/2020.04.03.023531

**Authors:** Sirjan Sapkota, Jon Lucas Boatwright, Kathleen Jordan, Richard Boyles, Stephen Kresovich

## Abstract

Cereal grains, primarily composed of starch, protein, and fat, are major source of staple for human and animal nutrition. Sorghum, a cereal crop, serves as a dietary staple for over half a billion people in the semi-arid tropics of Africa and South Asia. Genomic prediction has enabled plant breeders to estimate breeding values of unobserved genotypes and environments. Therefore, the use of genomic prediction will be extremely valuable for compositional traits for which phenotyping is labor-intensive and destructive for most accurate results. We studied the potential of Bayesian multi-output regressor stacking (BMORS) model in improving prediction performance over single trait single environment (STSE) models using a grain sorghum diversity panel (GSDP) and a biparental recombinant inbred lines (RILs) population. A total of five highly correlated grain composition traits: amylose, fat, gross energy, protein and starch, with genomic heritability ranging from 0.24 to 0.59 in the GSDP and 0.69 to 0.83 in the RILs were studied. Average prediction accuracies from the STSE model were within a range of 0.4 to 0.6 for all traits across both populations except amylose (0.25) in the GSDP. Prediction accuracy for BMORS increased by 41% and 32% on average over STSE in the GSDP and RILs, respectively. Predicting whole environments by training with remaining environments in BMORS yielded higher average prediction accuracy than from STSE model. Our results show regression stacking methods such as BMORS have potential to accurately predict unobserved individuals and environments, and implementation of such models can accelerate genetic gain.

## Introduction

Cereal grains provide more than half of the total human caloric consumption globally and amount to over 80% in some of the poorest nations of the world [1]. Sorghum [*Sorghum bicolor* (L.) Moench], a drought-tolerant cereal crop, is a dietary staple for over half a billion people of semi-arid tropics which is inhabited by some of the most food insecure and malnourished populations [2]. In industrialized countries, such as United States and Australia, grain sorghum is primarily grown for animal feed. But in recent years the uses of sorghum grain have expanded to baking, malting, brewing, and biofortification [3–5]. Therefore, genetic improvement of sorghum grain composition is crucial to mitigate the global malnutrition crisis, to increase efficiency of feed grains used in animal production, and to serve evolving niche markets for gluten-free grains.

In the last two decades, the use of genome-wide markers in prediction of genetic merit of individuals has revolutionized plant and animal breeding. Genomic prediction (GP) uses statistical models to estimate marker effects in a training population with phenotypic and genotypic data which is then used to predict breeding values of individuals solely from genetic markers [6, 7]. Training population size, genetic relatedness between individuals in training and testing population, marker density, span of linkage disequilibrium and genetic architecture of traits are some of the factors that can affect the predictive ability of the models [8–10]. Genomic prediction models are routinely studied and applied by breeding programs around the world in several crops. Novel statistical methods that are capable of incorporating pedigree, genomic, and environmental covariates into statistical-genetic prediction models have emerged as a result of extensive computational research [11].

One of the main advantages of GP is that breeders can use phenotypic values from some lines in some environments to make predictions of new lines and environments. Genomic best linear unbiased prediction (GBLUP) proposed by VanRaden [12] is probably the most widely used genomic prediction model in both plant and animal breeding. Since then GBLUP model has been extended to include G *×* E interactions resulting in improved prediction accuracy of unobserved lines in environments [13, 14]. Burguenño *et. al*. [13] found an increase in prediction ability of unobserved wheat genotypes by about 20% in multi-environment GBLUP model compared to single environment model. Also an extension of the GBLUP model, Jarquín *et. al*. [14] introduced a reaction norm model which introduces the main and interaction effects of markers and environmental covariates by using high-dimensional random variance-covariance structures of markers and environmental covariates. While most of the genomic prediction studies have been on individual traits, breeding programs use selection indices based on several traits to make breeding decisions. To address those challenges, expanded genomic prediction models that perform joint analysis of multiple traits have been studied using empirical and simulated data [15, 16]. Subsequent improvement in prediction accuracy from multi-trait model over single-trait model depends on trait heritability and correlation between the traits involved [15, 17].

Data generated in breeding programs span multiple environment and are recorded for multiple traits for each individual. While multi-environment models and multi-trait models are implemented separately, a single model to account for complexity of variance-covariance structure in a combined multi-trait multi-environment (MTME) model was lacking until Montesinos-López *et. al*. [18] developed a Bayesian whole genome prediction model to incorporate and analyze multiple traits and multiple environments simultaneously. Montesinos-López *et. al*. [18] also developed a computationally efficient Markov Chain Monte Carlo (MCMC) method that produces a full conditional distribution of the parameters leading to an exact Gibbs sampling for the posterior distribution. Another MTME model that employs a completely different method was proposed by Montesinos-López *et. al*. [19]. This method, called the Bayesian multi-output regression stacking (BMORS), is a Bayesian version of multi-target regressor stacking (MTRS) originally proposed by Spyromitros-Xioufis *et. al*. [20, 21]. This method consists of training in two stages: first training multiple learning algorithms for the same dataset and then subsequently combining the predictions to obtain the final predictions.

Genomic prediction for grain quality traits has previously been reported in crops such as wheat [22–24], rye [25], maize [26], and soybean [27]. Hayes *et. al*. [28] and Battenfield *et. al*. [23] used near-infrared derived phenotypes in genomic prediction of protein content and end-use quality in wheat. Multi-trait genomic prediction models can simultaneously improve grain yield and protein content despite being negatively correlated [24, 29]. In sorghum, grain macronutrients have shown to be inter-correlated among one another [30], which suggests the multi-trait models may increase predictive ability of individual grain quality traits. The ability to assess genetic merit of unobserved selection candidates across environments is promising for reducing evaluation cost and generation interval in the sorghum breeding pipeline where parental lines of commercial hybrids are currently selected on the basis of extensive progeny testing [31]. In order to extend capacities to performance index selection for multiple environments, we need to study and effectively implement MTME genomic prediction models in our breeding programs. In this study, we report the first implementation of genomic prediction for grain composition in sorghum, and the objective was to assess potential for improvement in prediction accuracy of multi-trait regressor stacking model over single trait model for five grain composition traits: amylose, fat, gross energy, protein and starch.

## Materials and methods

### Plant material

#### Grain sorghum diversity panel

A grain sorghum diversity panel (GSDP) of 389 diverse sorghum accessions was planted in randomized complete block design with two replications in 2013, 2014, and 2017 field seasons at the Clemson University Pee Dee Research and Education Center in Florence, SC. The GSDP included a total of 332 accessions from the original United States sorghum association panel (SAP) developed by Casa *et. al*. [32]. The details on experimental field design and agronomic practices are described in Boyles *et. al*. [33] and Sapkota *et. al*. [34]. Briefly, the experiments were planted in a two row plots each 6.1 m long, separated by row spacing of 0.762 m with an approximate planting density of 130,000 plants *ha*^*−*1^. Fields were irrigated only when signs of drought stress was seen across the field.

#### Recombinant inbred population

A biparental population of 191 recombinant inbred lines (RILs) segregating for grain quality traits was studied along with the GSDP. The parents of the RIL population were BTx642, a yellow-pericarp drought tolerant line, and BTxARG-1, a white pericarp waxy endosperm (low amylose) line. The population was planted in two replicated plots in randomized complete block design across two years (2014 and 2015) in Blackville, SC and College Station, TX. Field design and agronomic practices have previously been described in detail in Boyles *et. al*. [30].

### Phenotyping

The primary panicle of three plants selected from each plot were harvested at physiological maturity. The plants from beginning and end of the row were excluded to account for border effect. Panicles were air dried to a constant moisture (10-12%) and threshed. A 25g subsample of cleaned and homogenized grain ground to 1-mm particle size with a CT 193 Cyclotec Sample Mill (FOSS North America) was used in near-infrared spectroscopy (NIRS) for compositional analysis.

Grain composition traits such as total fat, gross energy, crude protein, and starch content can be measured using NIRS. Previous studies have shown high NIRS predictability of the traits used in feed analysis [35, 36]. We used a DA 7250™ NIR analyzer (Perten Instruments). The ground sample was packed in a gradually rotating Teflon dish positioned under the instrument’s light source and predicted phenotypic values was generated based on calibration curve for spectral measurements. The calibration curve was built using wet chemistry values from a subset of samples. The wet chemistry was performed by Dairyland Laboratories, Inc. (Arcadia, WI) and the Quality Assurance Laboratory at Murphy-Brown, LLC (Warsaw, NC). The details on the prediction curves and wet chemistry can be found in Boyles *et. al*. [30].

### Genotypic data

Genotyping-by-sequencing (GBS) was used for genetic characterization of the GSDP and RILs populations [30, 33, 37]. Sequenced reads were aligned to the BTx623 v3.1 reference assembly (phytozome) using Burrows-Wheeler aligner [38]. SNP calling, imputation and filtering was done using TASSEL 5.0 pipeline [39]. The TASSEL plugin FILLIN for GSDP and FSFHap for RILs population were used to impute for missing genotypes. Following imputation SNPs with minor allele frequency (MAF)*<*0.01, and sites missing in more than 10% and 30% of the genotypes in GSDP and RILs, respectively, were filtered. The number of genotypes studied for each population represent those with at least 70% of SNP sites. The genotype matrix with 224,007 SNPs from GSDP and 56,142 SNPs from RILs population was used for whole genome regression.

### Statistical analysis

The statistical software environment ‘R’ was used for model building and analysis [40]. The phenotypic values of the traits were adjusted for random effects of replications within environment using ‘lme4’ package in R [41]. Principal component analysis was done using the R package ‘factoextra’ [42]. Marker-based estimates of narrow sense (genomic) heritabilities were calculated using the SNP genotype matrix and phenotypic values using the R package ‘heritability’ [43]. A matrix with dummy variables ‘1’ and ‘0’ representing combinations of environmental variables (replication and year for GSDP, and replication, year and location for RILs) was used as co-variate in heritability estimation.

#### Single-trait single-environment (STSE) model

The following genomic best linear unbiased prediction (GBLUP) model was used to assess prediction performance of an individual trait from a single environment:

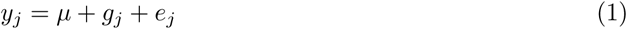

where *y*_*j*_ is a vector of adjusted phenotypic mean of the *j* th line (*j* = 1, 2,…, *J*). *µ* is the overall mean which is assigned a flat prior, *g*_*j*_ is a vector of random genomic effect of the *jth* line, with 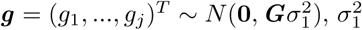 is a genomic variance, ***G*** is the genomic relationship matrix in the order *J × J* and is calculated [12] as 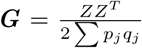, where *q*_*j*_ and *p*_*j*_ denote major and minor allele frequency of *j* th line respectively, and Z is the design matrix for markers of order *J × p* (*p* is total number of markers). Further, *e*_*j*_ is residual error assigned the normal distribution 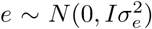 where *I* is identity matrix and *σ*^2^ is the residual variance with a scaled-inverse Chi-square density.

#### Bayesian multi-environment (BME) GBLUP model

Considering genotype *×* environment interaction can contribute to substantial amount of phenotypic variance in complex traits, we fit the following univariate linear mixed model to account for environmental effects in prediction performance:

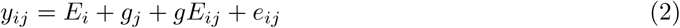

where *y*_*j*_ is a vector of adjusted phenotypic mean of the *j* th line in the *i* th environment (*i* = 1, 2,.., *I, j* = 1, 2,…, *J*). *E*_*i*_ represents the effect of *i* th environment and *g*_*j*_ represents the genomic effect of the *j* th line as described in equation 1. The term *gE*_*ij*_ represents random interaction between the genomic effect of *j* th line and the *i* th environment with 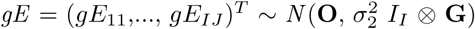, where 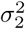 is an interaction variance, and *e*_*ij*_ is a random residual associated with the jth line in the ith environment distributed as 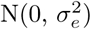 where 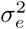 is the residual variance.

#### Bayesian multi-output regressor stacking (BMORS)

BMORS is the Bayesian version of multi-trait (or multi-target) regressor stacking method [19]. The multi-target regressor stacking (MTRS) was proposed by Spyromitros-Xioufis *et. al*. [20, 21] based on multi-labeled classification approach of Godbole and Sarawagi [44]. In BMORS or MTRS, the training is done in two stages. First, *L* univariate models are implemented using the multi-environment GBLUP model given in equation 2, then instead of using these models for prediction, MTRS performs the second stage of training using a second set of *L* meta-models for each of the *L* traits. The following model is used to implement each meta-model:

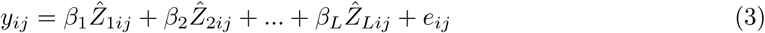

where the covariates 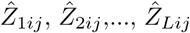 represent the scaled prediction from the first stage training with the GBLUP model for *L* traits, and *β*_1_, …, *β*_*L*_ are the regression coefficients for each covariate in the model. The scaling of each prediction was performed by subtracting its mean (*µ*_*lij*_) and dividing by its corresponding standard deviation (*σ*_*lij*_), that is, 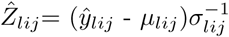, for each *l* = 1,…, *L*. The scaled predictions of its response variables yielded by the first-stage models as predictor information by the BMORS model. Simply put, the multi-trait regression stacking model is based on the idea that a second stage model is able to correct the predictions of a first-stage model using information about the predictions of other first-stage models [20, 21].

#### Performance of prediction model

All prediction models were fit using Bayesian approach in statistical program ‘R’. The STSE model (1) was fit using the R package ‘BGLR’ [45], BME model (2) and BMORS model (3) were fit using the R package ‘BMTME’ [19]. A minimum of 20,000 iterations with 10,000 burn-in steps was used for each Bayesian run.

The evaluation of prediction performance of models was done using a five-fold cross validation (CV), which means 80% of the samples were used as training set and testing was done on the remaining 20% for each cross-validation fold. The individuals were randomly assigned into five mutually exclusive folds. Four folds were used to train prediction models and to predict the genomic estimated breeding values (GEBVs) of the individuals in fifth fold (validation/test set). The accuracy of prediction for each fold was calculated as Pearson’s correlation coefficient (r) between predicted values and adjusted phenotypic means for the individuals in validation set. Each cross validation run, therefore, resulted in five estimates of prediction accuracy. The same set of individuals were assigned to training and validation across different traits and models tested by using *set*.*seed()* function in R. In order to avoid bias due to sampling, we performed 10 different cross-validation runs to calculate the mean and dispersion of the prediction accuracies.

## Results

### Phenotypic variation

A single calibration curve for NIRS was used for both populations studied. Table 1 outlines the summary statistics of NIRS predictions and phenotypic distribution and heritability of the grain composition traits. The cross validation accuracy (*R*^2^) of the NIRS calibration curve was moderately high to high, except for fat which had a moderate *R*^2^ value (0.41). We had a total of three environments (three years in one location) for the GSDP and four environments (two years in two locations) for the RILs. Traits were normally distributed except amylose in two 2014 environments in the RILs which had bimodal distribution (S1 Fig, S2 Fig). All traits showed significant variation in distribution across the environments, except starch in GSDP.

**Table 1.** Summary statistics of NIRS calibration and phenotypic distribution. *R*^2^ is the prediction accuracy and SECV is the standard error of cross validation for the NIRS calibration curve. Mean represents the phenotypic mean of the trait with its standard deviation (SD). *h*^2^ is the estimate of genomic heritability.

| Trait | NIRS |  | GSDP |  | RILs |  |
| --- | --- | --- | --- | --- | --- | --- |
| | $R^2$ | SECV | Mean ± SD | $h^2$ | Mean ± SD | $h^2$ |
| Amylose | 0.60 | 2.24 | 13.87 ± 2.98 | 0.24 | 11.49 ± 4.32 | 0.77 |
| Fat | 0.41 | 0.53 | 2.53 ± 0.57 | 0.54 | 3.07 ± 0.67 | 0.76 |
| Gross energy | 0.71 | 25.80 | 4108.33 ± 55.15 | 0.59 | 4124.56 ± 41.74 | 0.69 |
| Protein | 0.96 | 0.27 | 12.02 ± 1.45 | 0.39 | 11.43 ± 1.03 | 0.83 |
| Starch | 0.89 | 0.75 | 68.30 ± 2.44 | 0.44 | 68.37 ± 1.87 | 0.79 |

The genomic heritabilities of all traits except gross energy were significantly higher (p *<*0.05) in the RILs than in the GSDP (Table 1). Trait heritabilities were high in the RILs, with protein and gross energy having the highest and lowest heritabilities, respectively. In the GSDP, genomic heritability was moderately high for fat and gross energy, moderate for protein and starch, and low for amylose. The poor genomic heritability (0.24) of amylose in the GSDP was expected because only a very small proportion (1%) of accessions have low amylose as a result of *waxy* gene (Mendelian trait).

Fig 1 shows correlation between the adjusted phenotypic means for trait and environment combination. Starch was negatively correlated (p*<*0.001) with all other traits in both populations except for amylose in the RILs. Fat, protein and gross energy were significantly positively (p*<*0.001) correlated to each other across environments in both populations. The strongest positive correlation was between gross energy and fat, whereas the strongest negative correlations were found between starch *∼* gross energy and starch *∼* protein. Moderate to high positive correlation was observed between years for all traits (Fig 1). We conducted a principal component analysis (PCA) of correlation matrix for the traits in each environment. In both populations, the first component separated amylose and starch from the other three traits, whereas, the second component separated amylose from starch and gross energy from protein and fat (S3 Fig). The first component explained 78.8% and 75.9% of variation, and second component explained 6.3% and 9.8% of variation in the GSDP and RILs, respectively. The third principal component in the RILs separated proteins from fat and explained about 7.6% of the variation.

**Figure 1.**
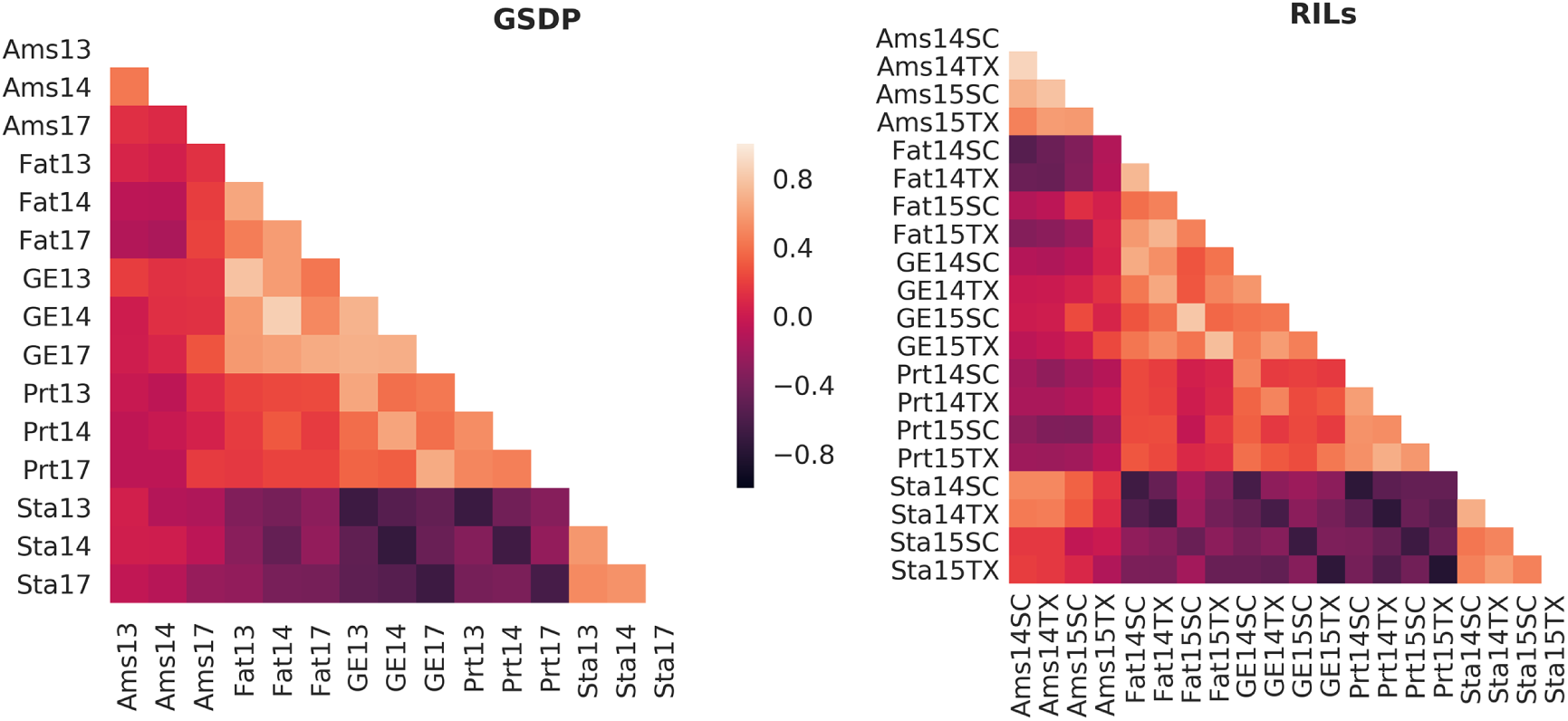
Correlation between traits across year and location combination for the two populations. Ams: amylose, GE: gross energy, Prt: protein, Sta: starch, SC: South Carolina, TX: Texas, and numbers in x and y-axes represent the year.

### Prediction performance in single and multiple environment

We first implemented GBLUP prediction model for single-trait single-environment (STSE). Prediction accuracies of the STSE model varied across environments in both populations (Fig 2). The environments 2014 in the GSDP and TX2014 in the RILs had highest average prediction accuracy but were not always the best predicted environment for all traits. While poorly predicted for amylose, the environments 2017 in the GSDP and TX2015 in the RILs had higher prediction accuracy for starch compared to all or most environments. Despite variation across environments and populations, the average prediction accuracies from the STSE were within the range of 0.4 to 0.6 for all traits except amylose (0.25) in the GSDP. The average prediction accuracy of the STSE model in the GSDP was positively correlated (r=0.86) with the genomic heritability of the traits. In the RILs, there was a positive correlation (r=0.77) between average prediction accuracy and genomic heritability for amylose, fat and gross energy, but the traits (protein and starch) with the highest heritabilities had relatively lower average prediction accuracies.

**Figure 2.**
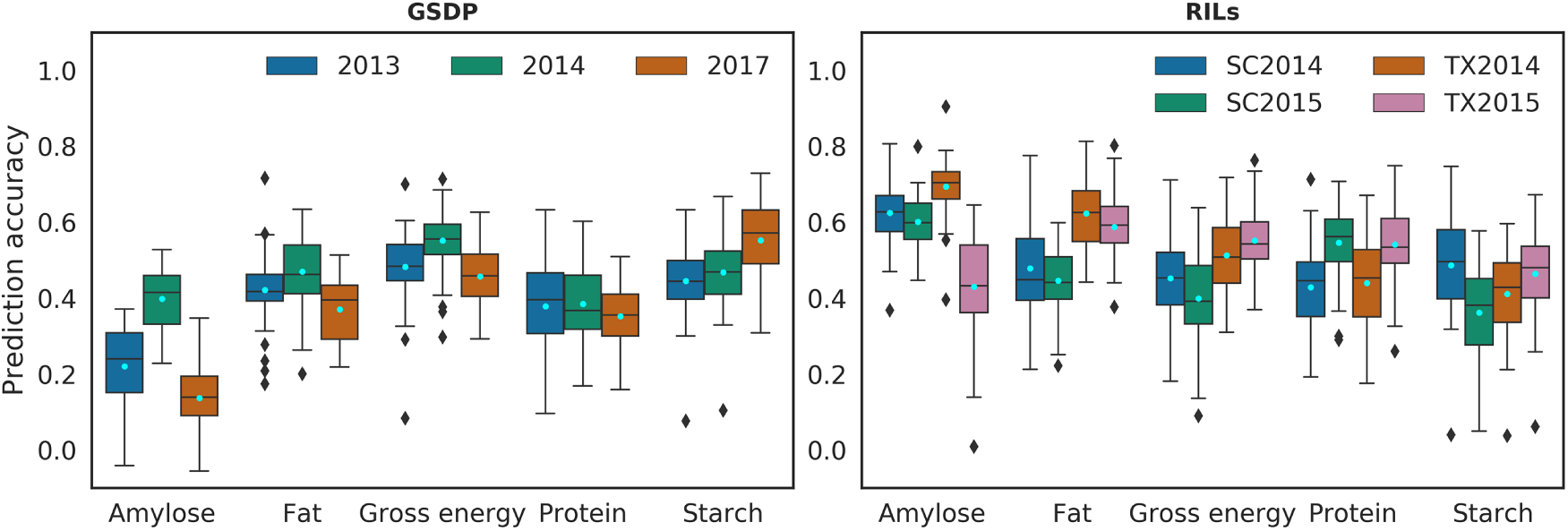
Prediction accuracy for single-trait single-environment model. Legend represents the environment/years. SC: South Carolina, TX: Texas. Pale blue dots represent the mean of prediction accuracy.

We didn’t see substantial improvement in multi-environment (BME) model over the STSE prediction accuracies (Fig 3). In the GSDP, the multi-environment models resulted in a decline in average prediction accuracy compared to the STSE model for fat (21%), amylose (10%) and protein (4%), however, no significant change was observed for gross energy and starch (S4 Fig, S1 Table). The prediction accuracy in the RILs increased by an average of 3% in the BME compared to the STSE, however, the overall trend of prediction accuracy for traits and environments remained unchanged (S4 Fig). The environment SC2014 showed consistent increase in accuracy for BME over STSE model across all traits with about 10% increase for protein (S2 Tab). Amylose in TX2015 environment had the single greatest increase (12%) in average prediction accuracy in the BME among all trait-environment combinations for the RILs.

**Figure 3.**
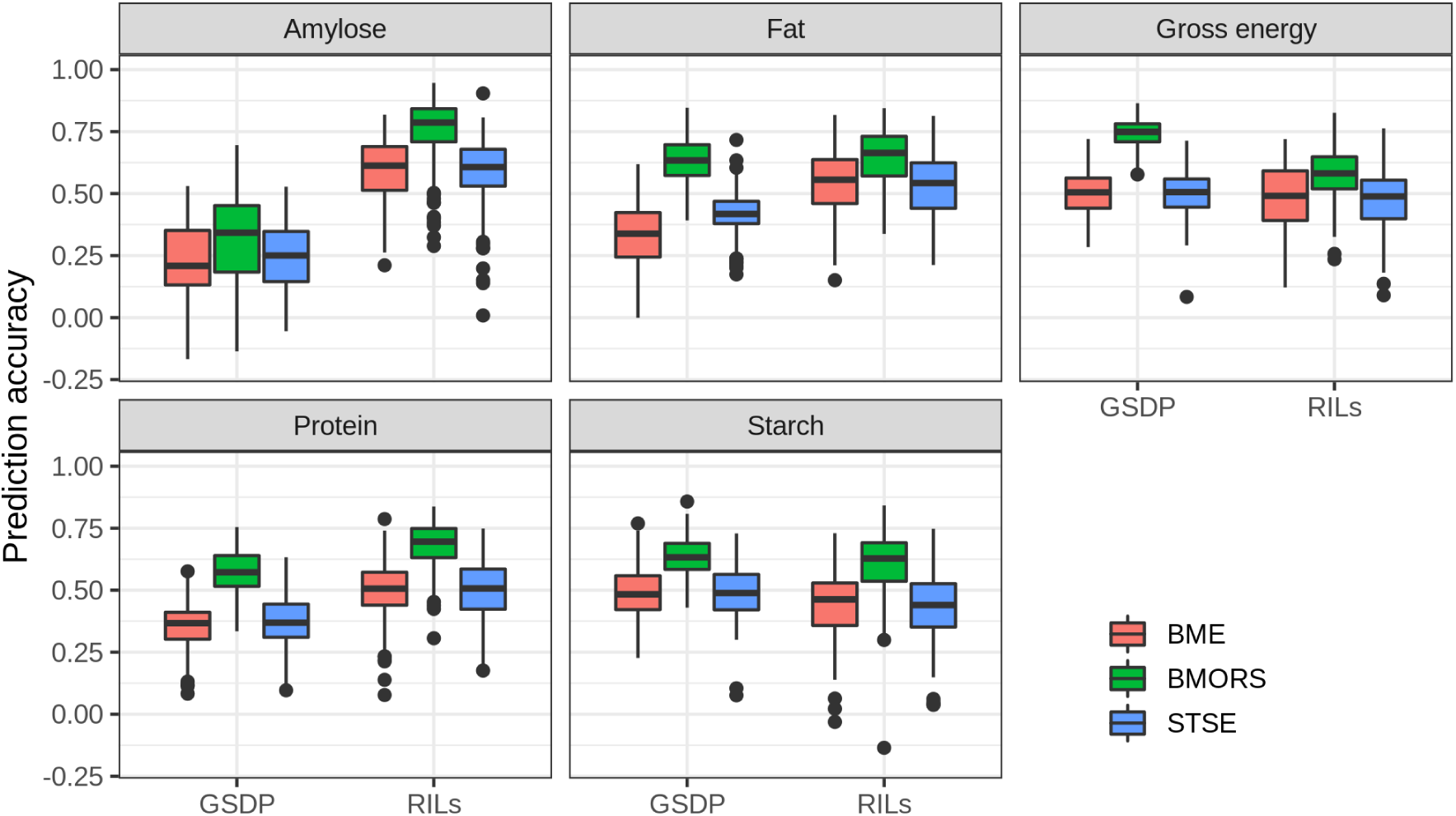
Average prediction accuracy of traits for the three prediction methods in the two populations.

### Bayesian multi-output regression stacking

We tested two different prediction schemes in the BMORS prediction model using the two functions *BMORS()* and *BMORS_Env()* as described in [19]. While the *BMORS()* function was used for a five-fold CV as described in the methods section, the *BMORS_Env()* was used to assess the prediction performance of whole environments while using the remaining environments as training.

#### Five-fold CV

The prediction accuracy from five-fold CV in BMORS increased by 41% and 32% on average over the STSE model in GSDP and RILs, respectively. Fig 4 shows the prediction accuracy of BMORS for each trait and environment combination across the two populations. While the percent change in accuracy varied across environments, the BMORS model nonetheless had higher average prediction accuracy than the STSE and BME models for all traits (Fig 3). The increase in average accuracy ranged from 11% (amylose, 2014) to 66% (amylose, 2013) in the GSDP with exception of amylose in 2017 (13% decrease), and 15% (fat, SC2015) to 60% (protein, TX2014) in the RILs (S1 Table). The increase in average prediction accuracy was higher (35%) for both locations in 2014 for the RILs, whereas, the year 2013 in the GSDP increased the most (S1 Table, S2 Table). Among the traits, protein (54%) had the greatest average increase in prediction accuracy in the GSDP, whereas in the RILs, protein and starch (42%) both showed the greatest increase.

**Figure 4.**
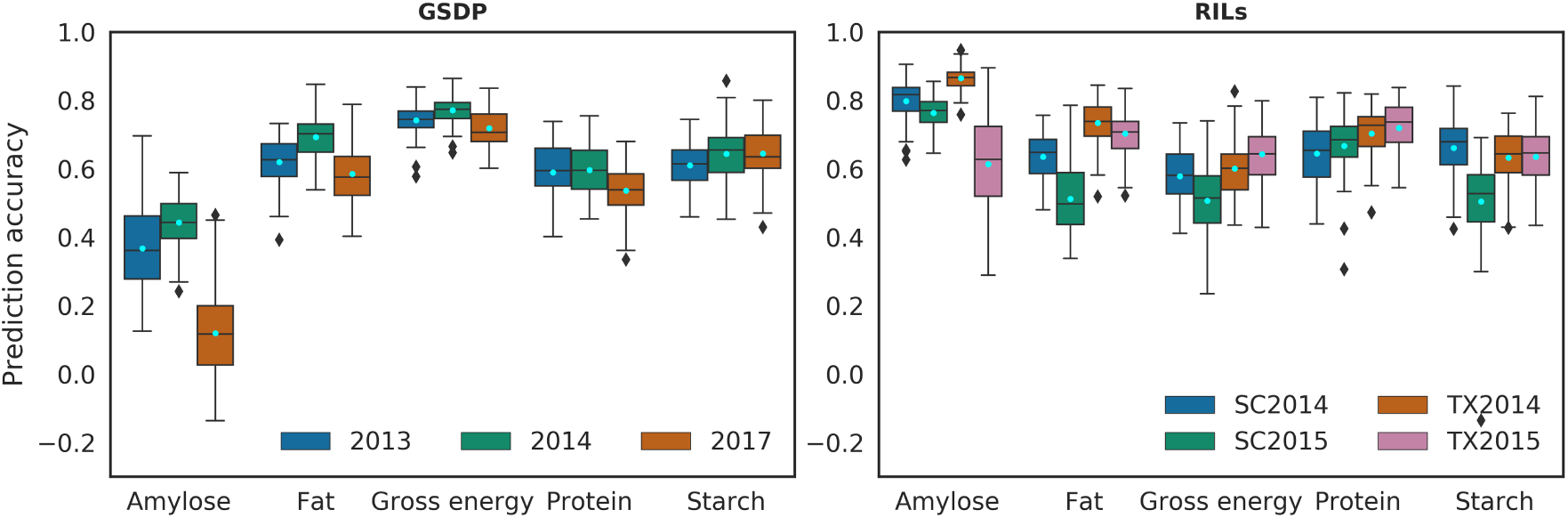
Prediction accuracy of BMORS model using five-fold CV. Legend represents the years/environment. SC: South Carolina, TX: Texas. Pale blue dots show mean of the prediction accuracy.

#### Prediction of whole environment

Predicting a whole environment using the BMORS model usually yielded higher accuracy than the mean prediction accuracy from the STSE model for each trait and environment combination (Fig 2, 5, Table 2). The distribution of prediction accuracy across trait and environment combination were, however, similar to the results from the STSE model. In the GSDP, little variation in prediction accuracies was observed across environments for gross energy, starch and protein, whereas, amylose and fat showed greater variability in prediction accuracy between environments. In the RILs, prediction accuracy for all traits except protein had high variability across the environments (Table 2).

**Table 2.**
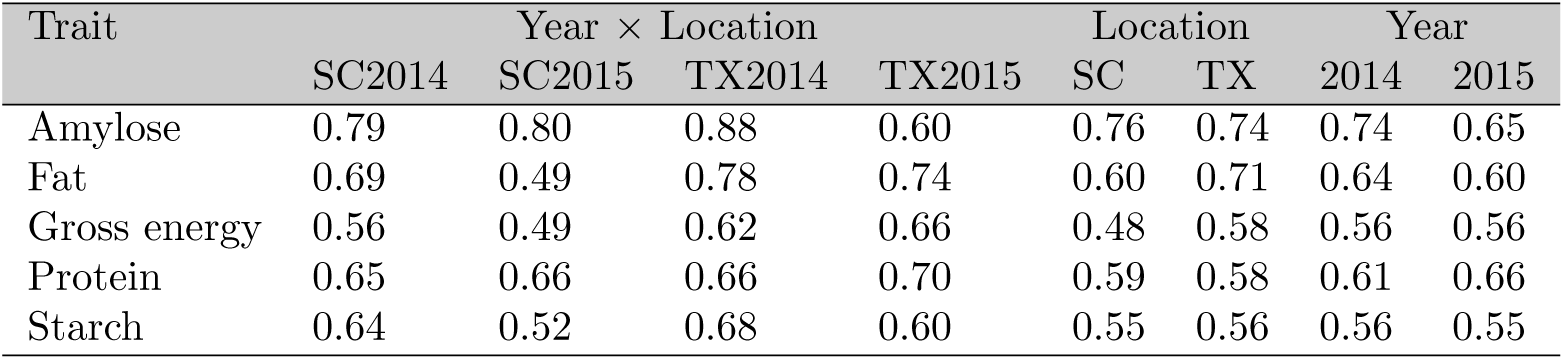
Prediction accuracy of the test environments predicted using the BMORS Env in the RILs.

**Figure 5.**
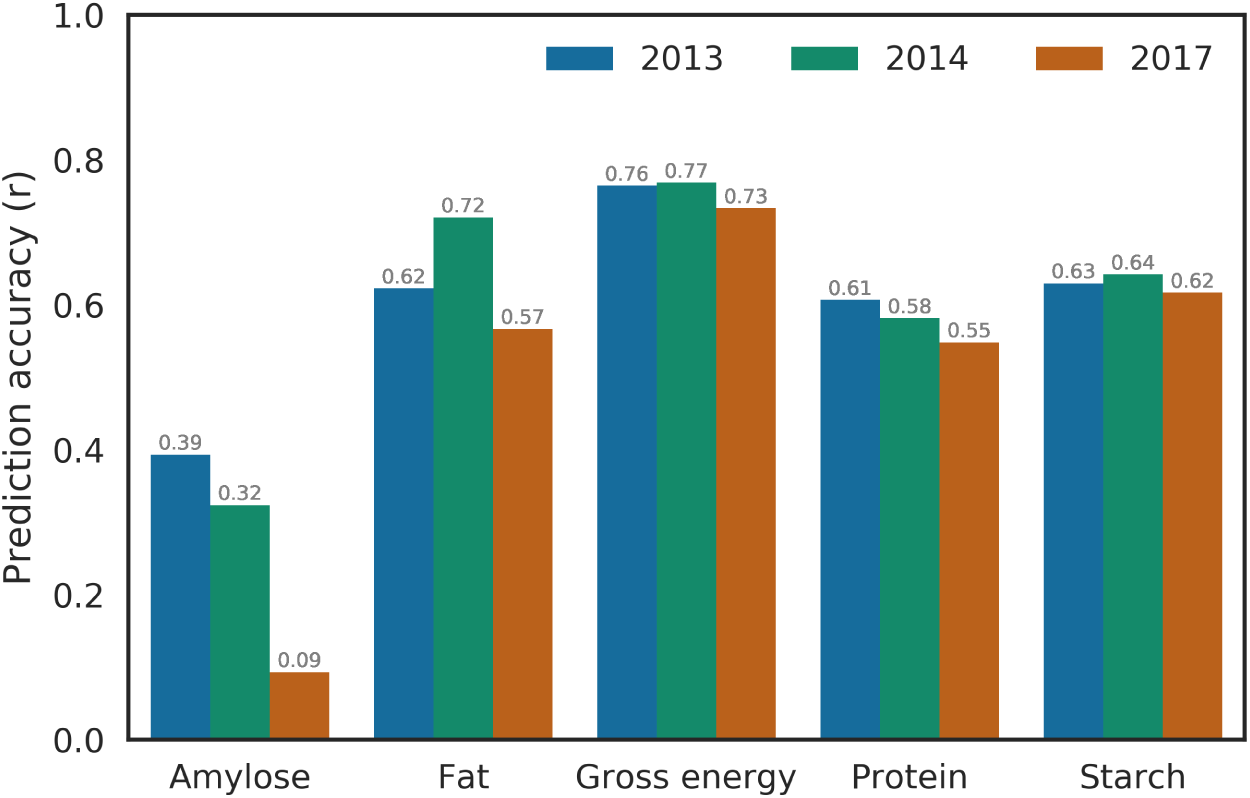
Prediction accuracy of the test environments predicted using the BMORS Env in the GSDP. Values on top of the bar represent the height of the bar.

In order to assess predictability by location or year in the RILs, we tested one location or year by training the BMORS model using the other location or year, respectively (Table 2). The Texas location had higher accuracy of prediction for fat (+0.11) and gross energy (+0.1) compared to South Carolina, but rest of the traits had similar prediction accuracy (difference *<*0.02). Prediction accuracy of whole years varied across traits, amylose (+0.09) and fat(+0.04) were higher in 2014, protein was higher (+0.05) in 2015, and starch and gross energy were similar.

## Discussion

Phenotyping for grain compositional traits is: 1) challenging and labor-intensive, 2) destructive for most accurate results, and 3) only performed after plants reach physiological maturity and are harvested. The use of genomic prediction for compositional traits will be extremely valuable because it increases selection intensity and decreases generational interval by overcoming the phenotyping challenges. Moreover, these traits are complex and quantitatively inherited so will benefit from genomic prediction’s ability to account for many small effect QTLs in estimating breeding values.

### Trait architecture and prediction accuracy

While the accuracy of NIRS calibration for traits in this study ranged from moderate to high, there was prediction error associated with NIRS prediction. However, it is unclear if and what effects NIRS prediction error had on genomic prediction. No direct correlation was observed between the genomic prediction accuracy and NIRS statistics for the traits studied. The trait with the lowest NIRS *R*^2^, fat, was predicted as well as or better than starch, protein and gross energy, which had NIRS *R*^2^ *>*0.7. Despite varying strength of correlations between traits across the two populations studied, the nature of relationship was similar for a given pair of traits, which is also in agreement with previous studies [30, 46, 47]. The strong negative relationship of starch and amylose to protein, fat and gross energy was further elucidated by the PCA analysis of phenotypic correlation matrix (S3 Fig). Since starch, protein and fat were measured on a percent dry matter basis, the strong correlation between them is expected.

Genetic relatedness and trait architecture are known to affect the accuracy of genomic prediction [8, 48]. The genetic relatedness between individuals and heritability of the traits were higher in the RILs than in GSDP (S3 Fig, Table 1). Those factors could be contributing to higher average prediction accuracy in the RILs. However, the average prediction accuracies for gross energy and starch were comparable between GSDP and RILs (Fig 3). Prediction accuracy in the GSDP could have been boosted by greater genetic diversity despite lower genetic relatedness [34]. Heffner *et. al*. [22] observed a prediction accuracy of 0.5-0.6 for wheat flour protein in two biparental populations. Guo *et. al*. [26] reported prediction accuracies of 0.44 and 0.8 for protein and amylose in rice diversity panel. Similar results were observed in our STSE models for protein content (Fig 2). Whereas, lower prediction accuracy of amylose in our diversity panel is probably due to the lack of sufficient low-amylose lines with the *waxy* gene [30]. While we couldn’t find any previous genomic prediction studies on starch, fat and gross energy, these traits are nutritionally one of the most important traits in cereal grains. The moderate to high prediction accuracy observed suggests implementation of genomic selection can improve genetic gain for these grain quality traits.

### Multi-trait regressor stacking

One of the daunting tasks of genomic prediction is estimating the effects of unobserved individuals and environments. As multiple traits are analyzed across several environments, the ability to combine information from multiple traits and environments can be crucial in increasing accuracy of prediction [13, 15, 16]. When the correlations among traits are high, prediction accuracies of complex traits can be increased by using multivariate model that takes this correlation into account [15, 18]. We fit a Bayesian multi-environment (BME) model (2) that takes the genotype *×* environment effects into consideration. In the GSDP, where environments were three years at the same location, the BME model showed a slight decline (7%) in average prediction accuracy which was mostly due to the two traits, amylose and fat (Supplementary Table S1). The RILs showed slight increase (2-3%) in prediction accuracy of traits when averaged over the environments, but there was variability across the environments (S2 Table).

We implemented two functions [*BMORS* () and *BMORS_Env* ()] which are not only used to evaluate prediction accuracy but are also computationally efficient [19]. The BMORS model (3) performs two-stage training by stacking the multi-environment models from all the traits. The five-fold cross validation conducted for BMORS was similar to the CV1 strategy of Montesinos-López *et. al*. [18]. The use of multi-trait models has been consistently shown to increase prediction accuracy over single-trait models across different crops and traits [15–17, 49]. The multi-target regressor stacking increased average prediction accuracy by 41% and 32% in the GSDP and RILs, respectively, as compared to the STSE prediction accuracy. Average prediction accuracy of all traits improved in BMORS over STSE and BME across both the populations (Fig 3). Consistent improvement in accuracy of BMORS is a result of the ability to use not only correlation between traits but also between environment in the model training [18,19]. The ability to accurately estimate genetic merit of lines in unobserved environments is of tremendous value in plant breeding. Our results show potential of *BMORS_Env()* function for predicting the whole environment. Testing a whole environment by training BMORS model using all other environments resulted in higher prediction accuracy for that trait-environment combination than using STSE or BME model. Prediction accuracy of all environments were 0.5 or higher with exception of amylose in GSDP, the reason for which we have discussed above (Fig 5, Table 2).

### Application for breeding

Grain quality traits such as starch and protein content have been under selection since the inception of phenotypic selection in modern breeding practices. More recently, total energy supplement of grain has gained attention for increasing feed efficiency in animal production, and a need exists for increasing total calories for human nutrition in the wake of global malnutrition crisis. Despite high correlations among these traits, the genetic variation underlying starch, protein and fat can be decoupled. [30] have shown major and minor effect QTLs underlying the three traits are distributed across the genome and are segregating in biparental populations. However, in practice, selection would be conducted simultaneously for these traits using a selection index rather than for individual traits. Velazco *et. al*. [31] observed an increase in predictive ability by using a multi-trait model for grain yield and stay green in sorghum, and argue that such an exercise would allow for using selection index for implementation of genomic selection for correlated traits. Increased prediction accuracy, improved selection index, and estimation of precise genetic, environmental and residual co-variances makes multi-trait multi-environment models preferable over univariate models [18]. The multi-trait regression stacking model we tested shows large scale improvement in model prediction and can be used in tandem with Bayesian multi-trait multi-environment (BMTME) model for parameter estimation and assessing prediction accuracy. The ability to estimate genetic effects and breeding values of unobserved environments will be of great advantage to predict performance in diverse environments and for implementation of selection theory.

## Conclusion

Phenotyping of grain compositional traits using near-infrared spectroscopy is labor-intensive, generally destructive, and time limiting. Therefore, the use of genomic selection for these traits will be extremely valuable. This study establishes the potential to improve genomics-assisted selection of grain composition traits by using multi-trait multi-environment model. The phenotypic measurements obtained from NIRS prediction were amenable to genomic selection as shown by moderate to high prediction accuracy for single trait prediction. While multi-environment model alone didn’t lead to much improvement over single environment model, stacking of regression from multiple traits showed substantial improvement in prediction accuracy. The prediction accuracy increased by 32% and 41% in the RILs and GSDP, respectively, when using the Bayesian multi-output regressor stacking (BMORS) model compared to a single trait single environment model. The ability to predict line performance in an unobserved environment is of great importance to breeding programs, and results show high accuracy for predicting whole environments using BMORS.

## Supporting information

**S1 Fig. Phenotypic distribution of grain composition traits in the RILs**. In the x-axes, SC: South Carolina, TX: Texas, numbers represent years. Values are percentage dry basis for protein, fat and starch; gross energy is in KCal/lb; and amylose is in percent of starch.

**S2 Fig. Phenotypic distribution of grain composition traits in the GSDP**. Numbers in x-axes represent years. Values are percentage dry basis for protein, fat and starch; gross energy is in Cal/g; and amylose is in percent of starch.

**S3 Fig. PCA analysis of correlation matrix between traits. a**. GSDP, and **b**. RILs. Ams: amylose, GE: gross energy, Prt: protein, Sta: starch, SC: South Carolina, TX: Texas. The numbers in the text represent years of the environment.

**S4 Fig. Prediction accuracy using five-fold CV in Bayesian multi-environment (BME) model. a**. GSDP, and **b**. RILs. Legend represents the environment/years. SC: South Carolina, TX: Texas. Pale blue dots represent the mean of prediction accuracy.

**S5 Fig. Heatmap for genomic relationship matrix calculated using vanRaden (2008). a**. GSDP, **b**. RILs. Trees show hierarchical clustering using Euclidean distance.

**S1 Table. Percent change in prediction accuracy over the single trait single environment model (STSE) model in the GSDP**. BME: Bayesian multi-environment, and BMORS: Bayesian multi-output regressor stacking.

**S2 Table. Percent change in prediction accuracy over the single trait single environment model (STSE) model in the RILs**. BME: Bayesian multi-environment, and BMORS: Bayesian multi-output regressor stacking.

## Acknowledgments

The authors would like to thank William L. Rooney and Brian K. Pfeiffer for their contributions to phenotyping of the recombinant inbred population at College Station, TX. Our appreciation goes to the Wade Stackhouse Fellowship, and Robert and Lois Coker Endowment for their support during the study. Clemson University’s computing cluster, Palmetto, was used for intensive data analyses. This study was funded by research grants from US Department of Energy ARPA-E TERRA (Award# DE-AR0000595, and DE-AR0001134).

## Data availability

The codes and data used in the study are available at github.com/sirjansapkota/GrainComp GS.

